# How do spike collisions affect spike sorting performance?

**DOI:** 10.1101/2021.11.29.470450

**Authors:** Samuel Garcia, Alessio P. Buccino, Pierre Yger

## Abstract

Recently, a new generation of devices have been developed to record neural activity simultaneously from hundreds of electrodes with a very high spatial density, both for *in vitro* and *in vivo* applications. While these advances enable to record from many more cells, they also dramatically increase the amount overlapping “synchronous” spikes (colliding in space and/or in time), challenging the already complicated process of *spike sorting* (i.e. extracting isolated single-neuron activity from extracellular signals). In this work, we used synthetic ground-truth recordings to quantitatively benchmark the performance of state-of-the-art spike sorters focusing specifically on spike collisions. Our results show that while modern template-matching based algorithms are more accurate than density-based approaches, all methods, to some extent, failed to detect synchronous spike events of neurons with similar extracellular signals. Interestingly, the performance of the sorters is not largely affected by the the spiking activity in the recordings, with respect to average firing rates and spike-train correlation levels.

## 1 Introduction

Accessing the activity of large ensemble of neurons is a crucial challenge in neuroscience. In recent years, Multi-Electrode Arrays (MEA) and large silicon probes have been developed to record simultaneously from hundreds of electrodes packed with a high spatial density, both *in vivo* [14, 2] and *in vitro* [10, 4]. With these devices, each electrode records the extracellular field in its vicinity and can detect the action potentials (or spikes) emitted by the neighboring neurons in the tissue. In contrast to intracellular recording, extracellular recordings do not give a direct and unambiguous access to single neuron activity and one needs to further process the recorded signals to extract the spikes emitted by the different cells around the electrodes. This complex problem of source separation is termed “spike sorting”. While various solutions for small number of channels (tens at max) can be found in the large literature on spike sorting algorithms [22], these new devices with thousands of channels challenge the *classical* approach to perform spike sorting.

Recently, a new generation of spike sorting algorithms have been developed to be able to deal with hundreds (or even thousands) of channels recorded simultaneously (see [16, 12] for recent reviews). The extent to which these modern spike sorting algorithm recover all the spikes from a neuronal population is still under investigations, and might differ depending on the species, tissue, cell types, activity level. While most of the real ground truth recordings [28, 19] are assessing the performance at the single cell level, in order to obtain an exhaustive assessment of the spike sorting performance at the population level, one must turn to use fully artificial or hybrid dataset [17, 6] to properly compare and quantify the performances of the algorithms. But even with such dataset, in most of the studies, errors are only measured as False Positive/Negative rates, and the reasons behind failures of the algorithms are often overlooked.

In this study, we focused on a key property of the spike trains, at the core of most of these failures, i.e. their fine temporal correlations. Indeed, temporal correlations are ubiquitous in the brain, and the higher the number of recorded cells because of the increased density of the probes, the more prominent they are. Correlations might have an important role in population coding ([3] for a review), but correlated activity for nearby cells results, in the extracellular signals, in overlapping activities and thus are harder to identify than isolated spikes. While pioneering work [21] claimed that template-matching based algorithms were more suited to recover overlapping spikes (either in space and/or time), the extent to which they are is not properly defined. In this work, our aim is to estimate how good (or bad) modern spike sorters are in recovering colliding spikes. What are the limits of the sorters, and what are the key parameters of the recordings and/or of the neurons that could influence these numbers?

## 2 Results

### 2.1 Simulated recordings

To test how robust the recently developed spike sorting pipelines are against spike collisions [28, 20, 8, 13, 15], we generated synthetic datasets using the `MEArec` simulator [6] (see Methods). More precisely, we took the layout of a NeuroNexus probe (A1×32-Poly3-5mm-25s-177-CM32 with 32 electrodes in three columns and hexagonal arrangement, a x- and y-pitch of 18 μm and 22 μm, respectively, and an electrode radius of 7.5 μm), and randomly positioned 20 cells in the vicinity of the probe (see Figure 1A), so that every simulated neuron has a unique *template* (i.e. average extracellular action potential). Figure 1B shows three sample templates with respectively low, almost null, and high similarity. The similarity between templates is computed as the cosine similarity of the flattened signals (see Methods) and the random generation of the positions and cell types of the simulated neurons (and thus of the templates) gives rise to the similarity matrix displayed in see Figure 1C. This similarity, as expected, decreases with the distance between the neurons, computed either from the ground-truth positions of the cells from the simulation or estimated as the barycenters of the templates (Figure 1D). The more negative the similarity is, the more templates are “in opposition”; the more positive it is, the more templates are “similar”. A similarity close to 0 means that templates do not overlap and are strongly orthogonal, i.e. dissimilar. Importantly, the simulations allowed us to cover rather uniformly the space of cosine similarities between templates, which will be used to assess the performance of spike sorters during collisions (Figure 1E).

**Figure 1:**
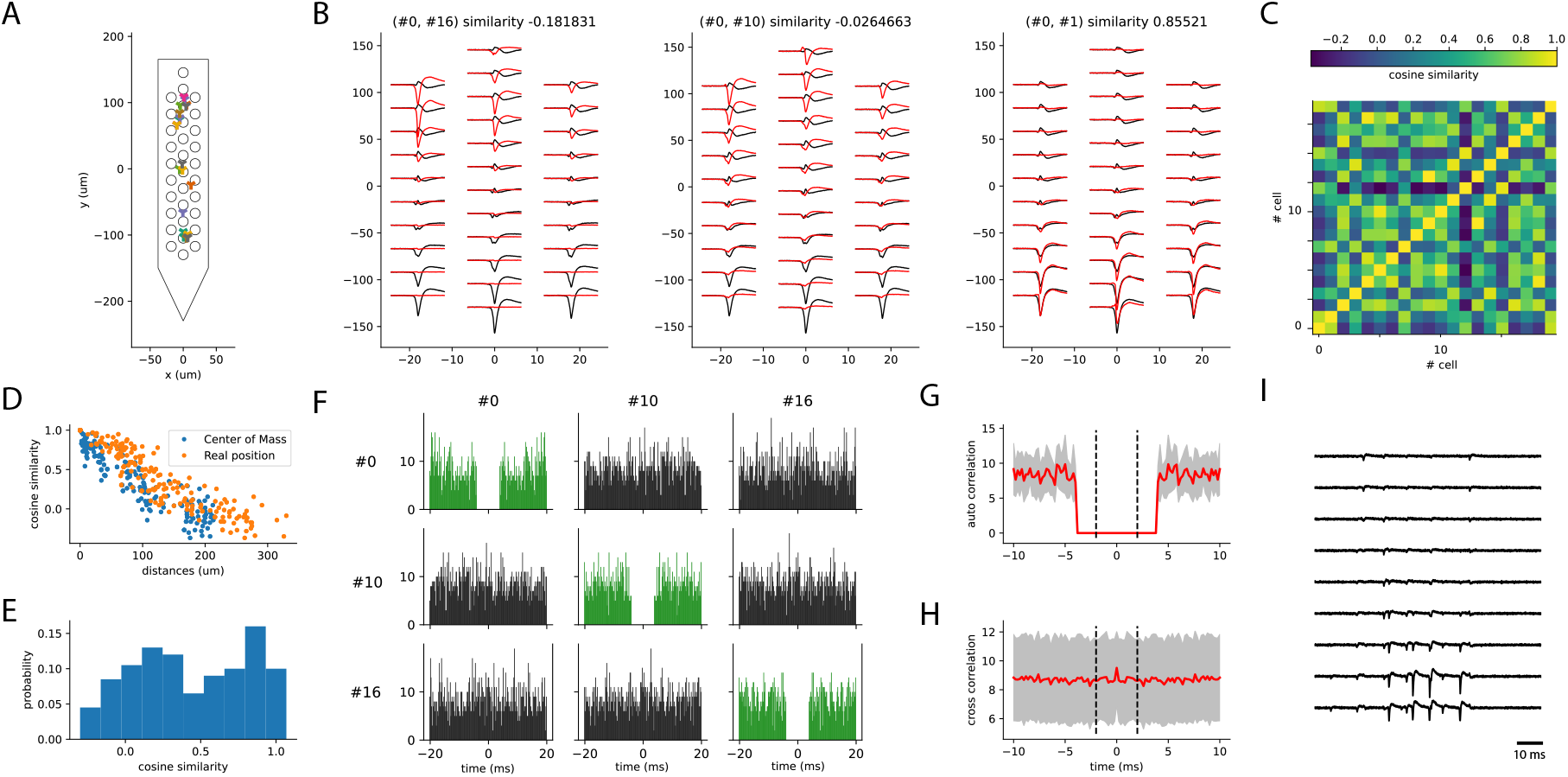
Generation of the synthetic recordings. **A)** 20 cells are randomly placed in front of a 32-channel NeuroNexus probe layout. The plot shows the location of each cell for one recording. **B)** Sample templates generated by neurons that are close too each other (#0 and #1) or far apart (#2). **C)** Cosine similarity matrix between all pairs of templates for a sample recording. **D)** Cosine similarity as function of the distance between the neurons, either using the real position from the simulations (orange circles), or the estimated barycenter of the templates (blue circles). **E)** Histogram of the cosine similarity distribution from one of the simulated recordings. **F)** Cross- and auto-correlograms for three sample spike trains. **G)** Average auto-correlograms of all units (red line, gray area represents the standard deviation). **H)** Average cross-correlogram over all pairs of neurons (red line, gray area represents the standard deviation around the mean). **I)** Sample traces from 10 channels of one synthetic recording.

To generate the spike trains, we first used a simple approach that forced all the neurons to fire as independent Poisson sources at a fixed and homogeneous firing rate of 5 Hz. To make the simulation more biologically plausible, we pruned all spikes breaking a refractory period violation of 4 ms. The resulting auto- and cross-correlograms for three sample units are shown in Figure 1F (auto-correlograms are in green on the diagonal), while Figure 1G and H display the average (red line) and standard deviation (grey shaded area) auto- and cross-correlation among all units, respectively. A sample snippet of the generated traces from one recording is shown in Figure 1I, for a subset of 10 channels out of 32. Due to the independence of the Poisson sources, both the average cross-correlograms (Figure 1G) and auto-correlograms – outside the ±4 ms used as refractory period – (Figure 1H) are flat.

### 2.2 Global performance of the spike sorters

In order to assess the global performances of the sorting procedure on our synthetic datasets, we generated 5 recordings with various random seeds and averaged the results. Figure 2 summarizes the main findings. First, we noticed that, as seen in Figure 2A, the run time was roughly constant across sorters, except for HDSort, with its higher run time. The number of well detected units is similar among sorters, as shown in Figure 2B, but it is worthwhile noticing that Kilosort 3 is the only sorter producing many false positive and redundant units (see Methods for classification of units). Kilosort 2 and HDSort also identify more false positive then well detected units. Importantly, we did not perform any curation of the spike sorting output, but we consider the raw output of each sorter as is.

**Figure 2:**
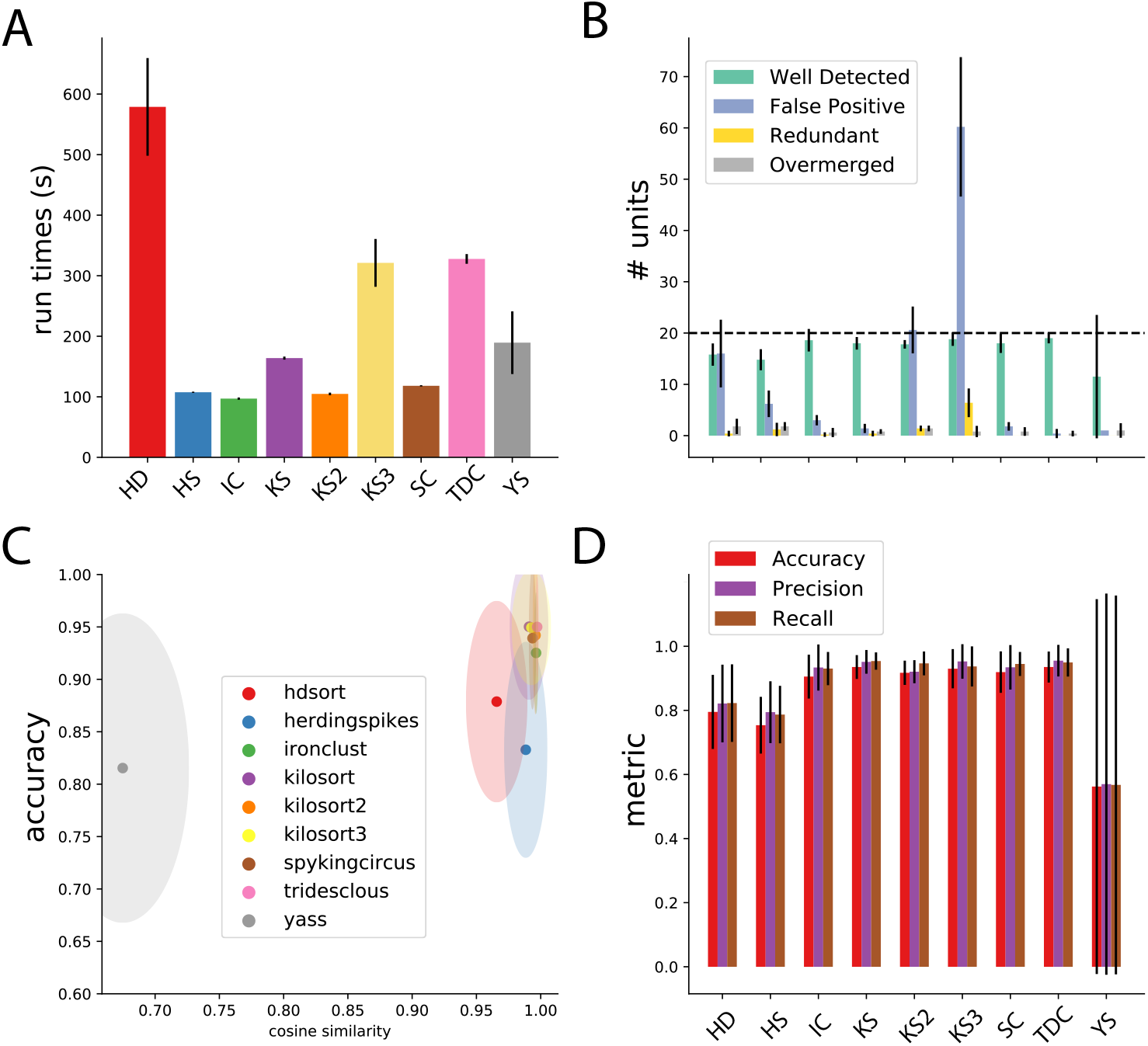
Spike sorting performance. **A)** Average run times over 5 different recordings (see Methods) for all the tested sorters. Errors bars indicate the standard deviation over multiple recordings. **B)** Average number of cells found by the sorters that are either well detected, redundant, overmerged or considered as false positive (see Methods). Error bars indicates standard deviation over multiple recordings. **C)** The average cosine similarity between templates found by the sorters and ground-truth templates, as function of the accuracy for the given neurons. Ellipses shows standard error of the means in cosine similarity (x-axis) and accuracy (y-axis). **D)** Average metrics (accuracy, precision, recall, see Methods) for all the sorters. Error bars show standard deviation over multiple recordings.

To check whether all sorters correctly *discovered* all templates, we computed the cosine similarity between the ground-truth templates from the simulations and the ones found by the sorters, comparing such a metric with the accuracy. As it can be seen in Figure 2C, all sorters are on average finding the correct templates, with the notable exception of YASS (in grey) and to some less extent HDsort (in red). Nevertheless, the overall accuracy of most of the spike sorters is relatively high (~0.95), except for HDsort and Herdingspikes which yield lower scores (Figure 2D). However, this averaged number does not tell us anything regarding the nature of these errors. While this error rate might seem low, it is likely that it is crucial, since it can potentially originate from the collisions, and thus from the correlations among neurons.

### 2.3 Spike sorting performance is affected by spike collisions

Using fully synthetic recordings with exhaustive ground truth, we can look at how good individual spike sorters perform specifically with respect to spatio-temporal collisions. To do so, we computed the *collision recall* (see Methods) as a function of the lag between two spikes, for a given pair of neurons. By averaging over multiple pairs of ground-truth neurons with similar template similarity (and over multiple recordings, see Methods), we can obtain a picture of how accurate the sorters are specifically with respect to the spike time lags and the similarities between templates. Figure 3 displays the collision recall per sorter as a function of the lag (x-axis), colored by the similarity between templates. Each panel shows the performance of a different spike sorter. One can immediately see that only few sorters are able to accurately resolve lag correlations that are close to zero, even when templates are strongly orthogonal (low cosine similarity). This is the case for Kilosort 1 and 2, and for Spyking-circus, all of which use a template-matching procedure that should theoretically explain this behavior. However, while performances are still good for Kilosort 1 and 2 even when the average cosine similarity between pairs is increased, they slightly degrade for Spyking-circus. Density-based sorters (HerdingSpikes and Ironclust), on the other hand, do not have a spike collision resolution strategy and this is reflected by their overall poorer performance. It is interesting to notice that Tridesclous, HDSort, YASS, and Kilsort 3, also using a template-matching based procedure to resolve the spikes, are not properly resolving the temporal correlations even for dissimilar templates. Different template-matching strategies are probably the cause of the differences among sorters. For example, HDSort and HerdingSpikes do not implement any strategy for spike collision resolution [9] and that is reflected in the quick degradation of performance as template similarity increases. KiloSort 1 and 2 used a GPU-based implementation of the k-SVD algorithm [1], used in matching learning as a dictionary learning algorithm for creating a dictionary for sparse representations. By doing so, it performs a reconstruction of the extra-cellular traces via orthogonal template matching pursuit, which is an enhancement of the greedy template matching pursuit (used in Spyking-circus and Tridesclous) more robust when templates are non-orthogonal. This might explain the boost in performance especially striking for templates with high similarity (*similarity* > 0.8).

**Figure 3:**
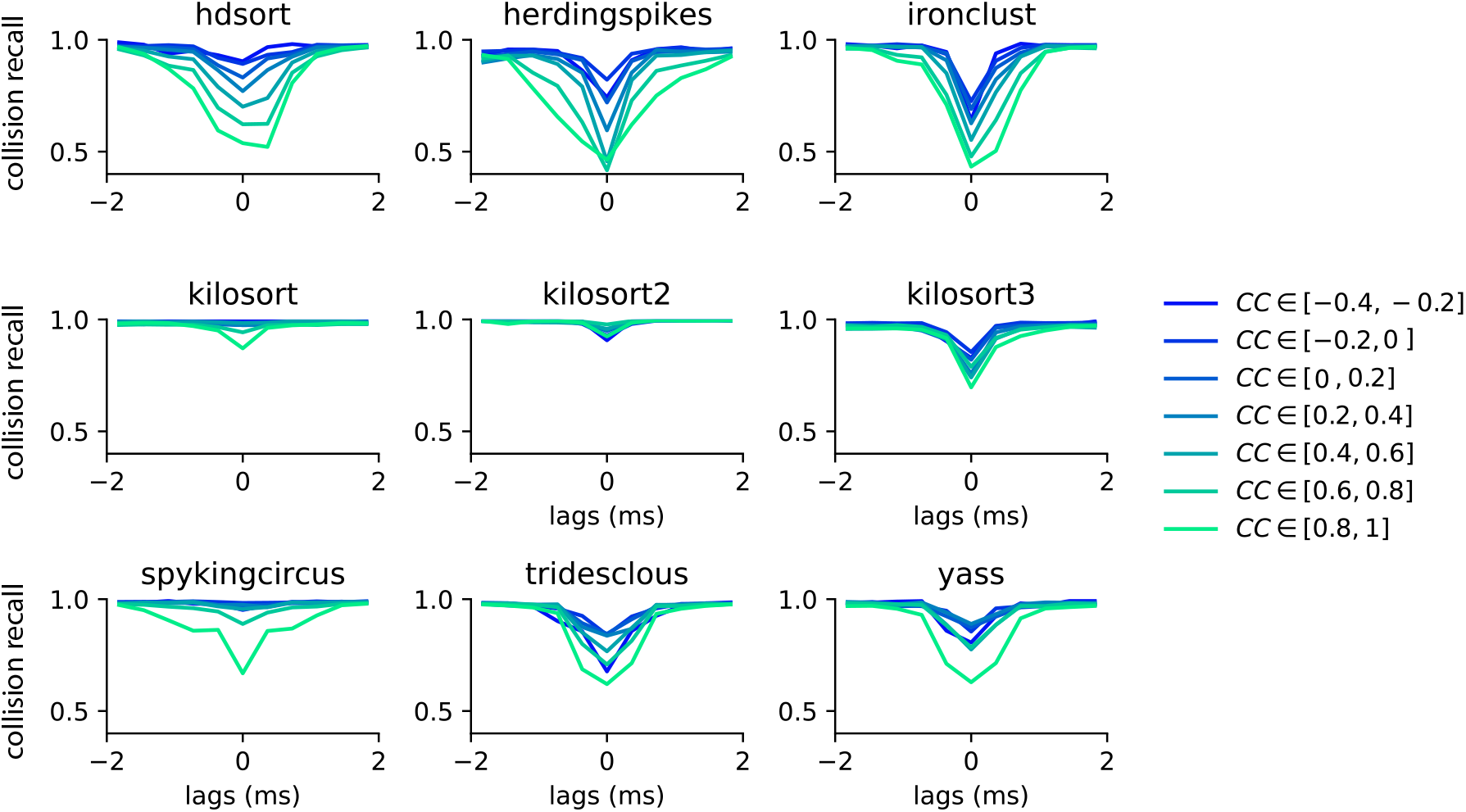
Collision recall per sorter. Error (quantified as the collision recall, see Methods) for various sorters and for all possible lags (between −2 and 2 ms), as function of the similarity between the pairs of templates (color code). All curves are averaged over multiple pairs and multiple recordings (see Methods).

### 2.4 Generation of controlled spike collision simulated data

The results shown in the previous section have been obtained only in a particular regime of activity, with all neurons firing independently as Poisson sources with an average firing rate of 5 Hz. However, neurons usually do not fire independently of each other, but rather have intrinsic correlations, also depending on different brain areas, brain states, and species. In addition, the average firing rates can also largely vary depending on brain areas. As an example, it is well known that Purkinje cells in the cerebellum have a very high firing rate [24] that networks tends to synchronize their activity either in slow waves during sleep [25], or during pathological activity (such as epileptic seizures [26]). Therefore, assessing how performances may vary during different conditions is important to generalize our observations.

In order to study how spike sorting is affected by correlations and firing rates, we used a mixture procedure [5] that allowed us to control precisely the shape of the auto- and cross-correlograms for the injected spike trains. More precisely, we decided to explore in a systematic manner three rate levels (5, 10 and 15 Hz), and three correlation levels (0, 10, and 20 %). Note that the 5 Hz firing rate with 0 % correlation corresponds to the scenario displayed in Figures 2–3.

Figure 4 shows the average of cross- and auto-correlograms and the spike trains of a recording where cells are firing as independent Poisson sources at 5 Hz in panels A-C (and thus with 0 % correlation, as shown by the flat average cross-correlograms in Figure 4A) and at 15 Hz with 20 % correlation (Figure 4D-F). Even though experimental recordings would contain a broader spectrum of firing rates and correlations, here we focus on assessing how different firing regimes affect spike sorting performance in a controlled setting. One would expect that the increased density of spikes (both in terms of firing rates and of synchrony) should degrade the performance of the spike sorters by affecting both the clustering step and the template-matching step, which in turn would degrade the resolution of spike collisions.

**Figure 4:**
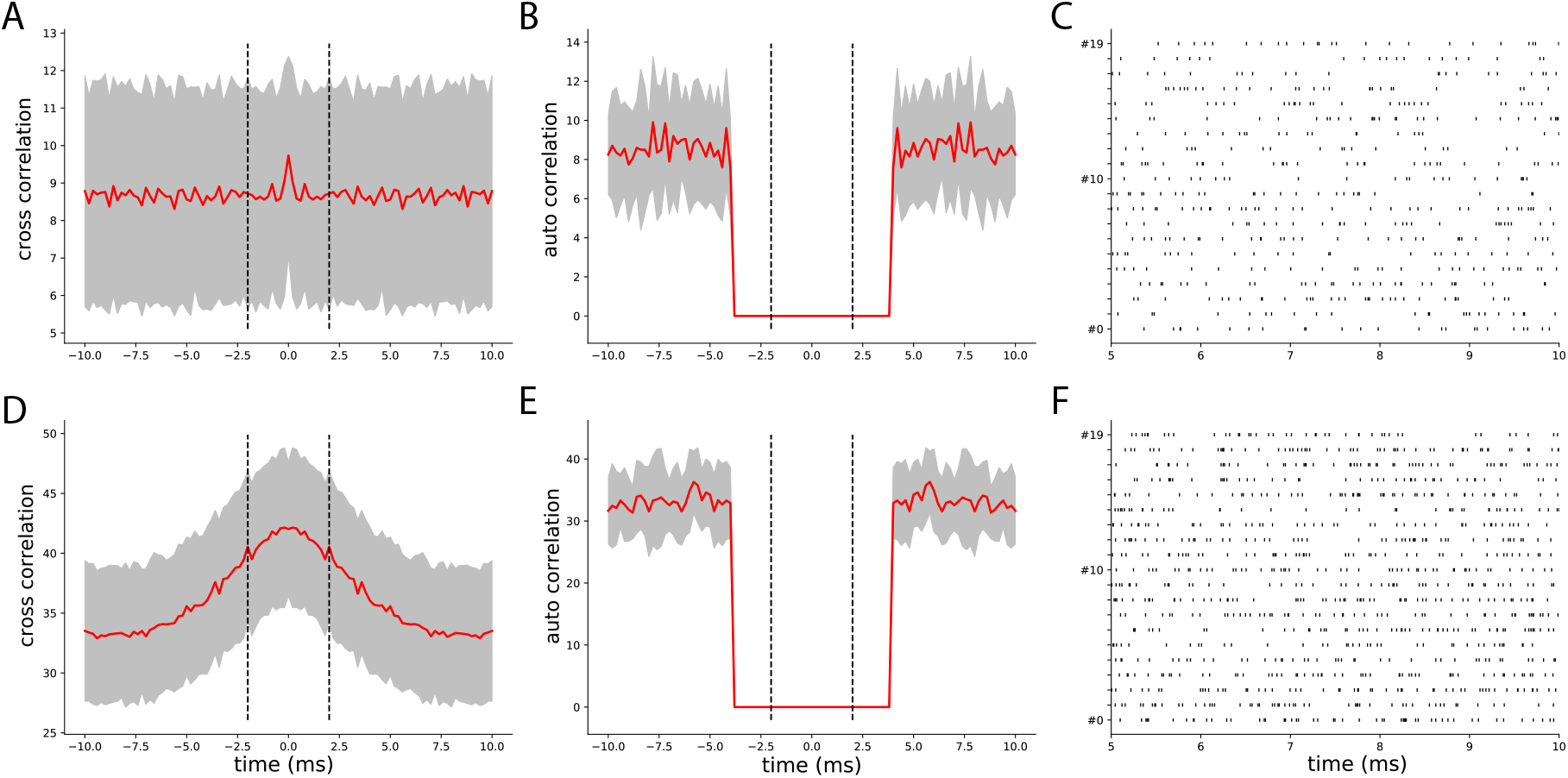
Controlling spike trains correlations and firing rates. **A)** Average cross-correlograms between all pairs of distinct neurons firing as independent Poisson sources at 5Hz (red curve, gray area represents the standard deviation) **B)** Same as **A**, but for auto-correlograms. **C)** Rater plot showing the activity of the uncorrelated neurons firing at 5Hz. **D-F** Same as **A-C**, but for a rate of 15 Hz and 20 % correlation.

### 2.5 Do correlations and firing rates affect spike sorting of spike collisions?

To assess whether firing rate and spike train correlation affect spike sorting performance, we selected all unit pairs with a similarity greater than 0.5. We first averaged the recall curves for all template similarities (i.e. we averaged the curves with similarity greater than 0.5 shown in Figure 3). In Figure 5A we show the recall with respect to the spike lags averaged over all 9 configurations (3 firing rates x 3 correlations) for each sorter. The thick line represents the mean recall and the shaded area is the standard deviation over different rate-correlation configuration. All sorters, except YASS, appear to have a very consistent curve (low standard deviation) over different configurations and do not seem affected by changes in average firing rates and correlations in the spike trains. YASS’ large standard deviation can be explained by looking at individual recall curves at different rate-correlation regimes (Figure 5 - figure supplement 1 - yellow lines): the spike sorting performance degrades with increasing firing rates, but it does not seem to be strongly affected by increased correlation rates. However, we should stress that since the collision recall is a relative measure, the same value for a larger number of spikes (when firing rate is increased) means that overall, there are more misses for all sorters.

**Figure 5:**
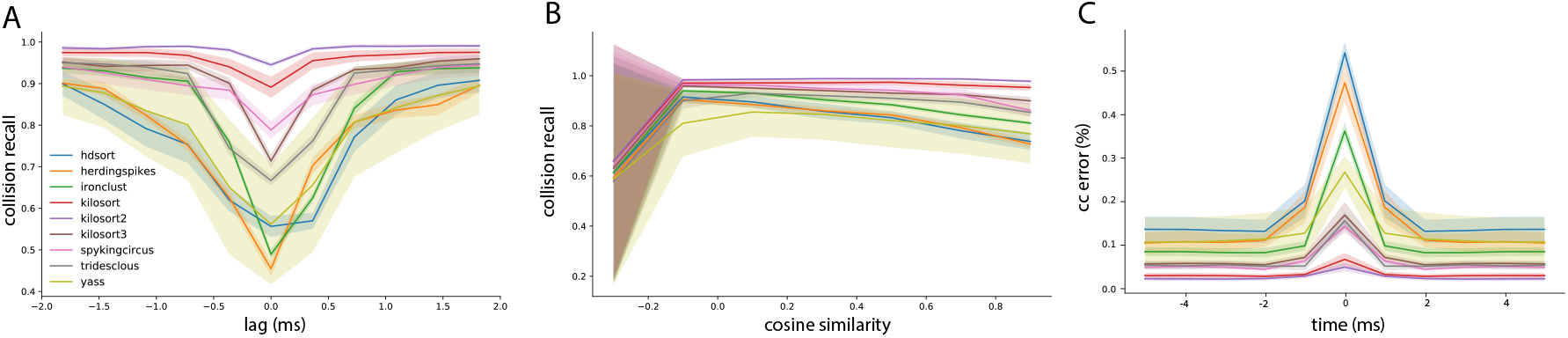
Spike sorting performance for different conditions. **A)** Average collision recall over the 9 conditions shown in Figure 5 - figure supplement 1 (3 firing rate levels and 3 correlation levels) as function of the lag between spikes, for pairs of cells with cosine similarity higher than 0.5. The shaded area shows the standard deviation over the conditions. **B)** Similarly as **A**, the average collision recall as function of the cosine similarity between pairs of cells. **C)** Mean relative error between the ground-truth cross-correlograms and the estimated ones, for all sorters, averaged over all pairs with a similarity higher than 0.5

Similar considerations can be done by looking at the average recall with respect to template similarity (Figure 5B). To construct this plots, we integrated the curves in Figure 3 over lags for different cosine similarities. Also in this case, the curves appear consistent (low standard deviation) with the exception of YASS, for which recall is reduced with increased firing rate regimes (Figure 5 - figure supplement 2 - yellow lines). It is worth noticing that when the cosine similarity becomes negative, all the sorters perform very poorly in properly resolving the overlaps. This could be explained by the fact that when a pair of templates is anti-parallel (for example in the left panel of Figure 1A), a subset of electrodes might show a negative signal for one template and a positive signal from the other (due to return currents in the dendritic signals [11]). Effectively, when a spike collision between the two spikes occur, this would lower the amplitude of the negative peak, which could reduce the detectability of the spike.

The collision recall metric is mostly useful to obtain a quantitative insight on the behavior of the spike sorting algorithms, but how do these errors transpose in practical situations? To assess this, we measure the relative error (in percentage) between the ground-truth cross-correlograms and the ones computed from the spike sorting outputs. We then averaged these error curves among all recordings and experimental conditions (firing rates and synchrony levels). As shown in Figure 5, the error in the estimated cross-correlogram can be as large as more than 50% for small lags, and for some spike sorting algorithms such as HDsort, HerdingSpikes or IronClust. Moreover, it is also worth noticing the baseline error rate is not the uniform across sorters. From this metric, we can again conclude that template-matching based spike sorting algorithms such as KiloSort (1, 2, and 3), Spyking-circus or Tridesclous are much better to resolve fine temporal correlations among neurons.

## 3 Discussion

In this study, we showed in a systematic and quantitative manner how spatio-temporal correlations can be underestimated during spike sorting. Using synthetic datasets, we compared a large diversity of modern spike sorters and showed how they behaved as function of the similarity between the templates and the temporal lags between spikes. As expected, the closer the spikes are in time, the harder is it, for all sorter, to properly resolve the overlaps. However, more interestingly, the more similar the templates are, the higher the failures are. These failures are striking especially for spike sorters that are not relying on template-matching based approaches (HerdingSpikes, Ironclust). For the ones using a template-matching based approach (KiloSort, Spyking-circus, Tridesclous, HDSort), the problem is less pronounced (with the exception of HDSort) but still present, and therefore this phenomenon should be taken into account when making claims about the synchrony.

**Figure 5 - figure supplement 1:**
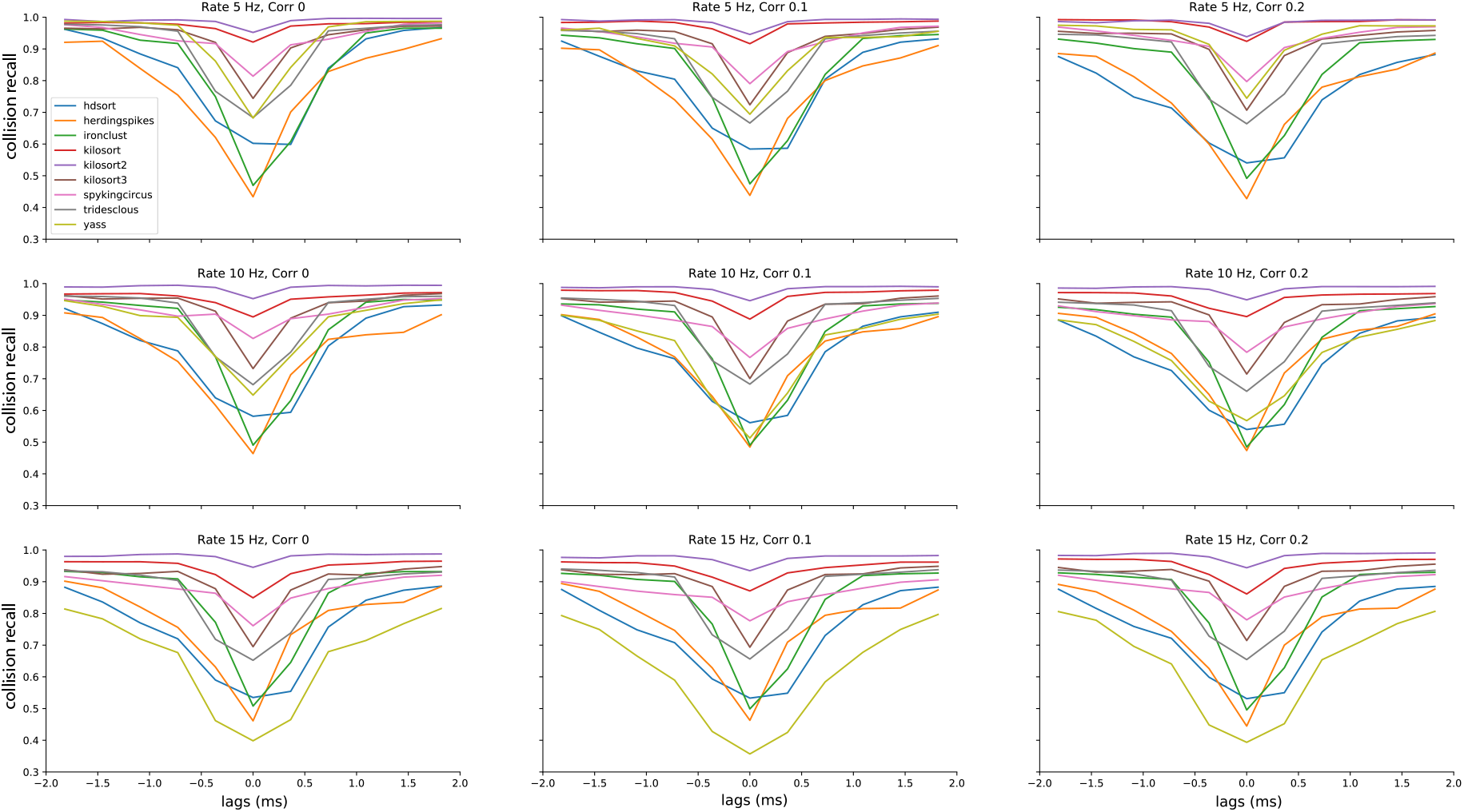
Average performances of the spike sorters as function of the temporal lags. Each panel shows the average collision recall for template pairs with a similarity above 0.5 for a different condition, in terms of firing rate and correlation levels.

**Figure 5 - figure supplement 2:**
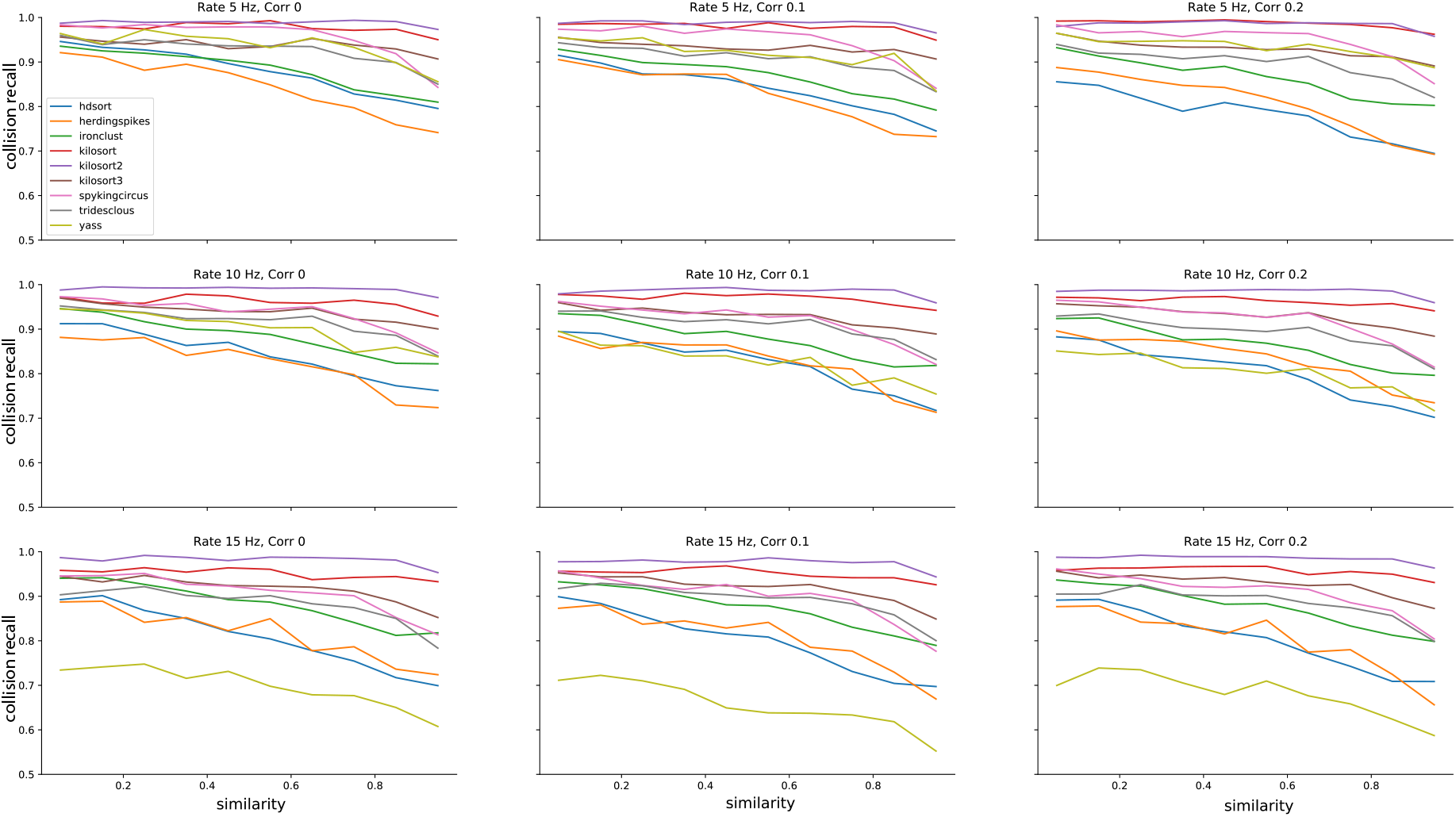
Average performances of the spike sorters as function of the template similarity. Each panel shows the average collision recall over all lags in [−2, 2] ms for a different condition, in terms of firing rate and correlation levels.

To our surprise, the global behavior of the spike sorters did not depend much on the overall firing rate and/or the correlation levels. This allows us to generalize the findings and we think that the quantitative results shown here could be translated to various *in vitro* or *in vivo* recordings from different brain regions and species. As shown in Figure 5, while the variability over different conditions is rather high for some algorithms, template-matching based algorithms tend to be rather robust and overall better in resolving spike collisions. This is a very encouraging sign towards a unified and reproducible automated solution for spike sorting [17, 7], agnostic of the recording conditions.

The results shown in the paper were obtained with purely artificial recordings, since we need exhaustive information on the ground-truth spiking activity of all neurons to quantitatively compare and benchmark different spike sorters. However, it would be interesting to generalize these observations with real recordings, assuming one would have a proper ground truth at the population level. Indeed, such a ground truth is needed to compute the *collision recall* and see how sorters behave as function of lags and similarities between templates. To our knowledge, such a ground truth does not exists [28, 19, 9]. While one could try to generate an “approximated” ground truth by combining the output of several spike sorters with an *ensemble* spike sorting approach (as in [7]), the disagreements among sorters are currently so high that this process is hard if not impossible, if one want to sample from a large number of pairs.

While missing spikes for very dissimilar templates and small lags is problematic, the errors made for very similar templates may be less frequent depending on the probe layout and neuronal preparation. Indeed, such errors strongly depends on the distribution of template similarities between all pairs of recorded cells, and this distribution might differ from recording to recording. For example, in the retina [27] one would expect highly synchronous cells, of the same functional type, to be far apart from each other because of an intrinsic tiling of the visual space. Such properties are unknown *in vivo* or in cortical structures, but might bias the distribution of template similarities between nearby neurons, and thus modify the estimation of collision recalls.

## 4 Methods

All the code used to generate the figures is available at https://spikeinterface.github.io/.

### 4.1 Simulated datasets

We used the MEArec simulator [6] to generate synthetic ground truth recordings. In brief, MEArec uses biophysically detailed multicompartment models to simulate the extracellular action potentials, or so called “templates”. For this study, we used 13 cell models from layer 5 of a juvenile rat somatosensory cortex [18, 23]. Templates are then combined with spike trains and slightly modulated in amplitude to add physiological variability. Additive uncorrelated Gaussian noise is finally added to the traces. We generated simulated recordings with 20 neurons randomly positioned in front of the probe, a noise level of 5 μV and a sampling rate of 32 kHz. To obtain more robust results, we generated 5 recording per conditions with various random seeds. The spike times were kept unchanged, but the positions and the templates of the 20 neurons were changed in each of the individual recording. This allowed us to populate the distribution of cosine similarities between pairs.

### 4.2 Generating spike trains with controlled correlations

To generate the recordings with various firing rates and correlations levels, we used the mixture process method described in [5]. Since by default the method generates controlled cross-correlograms with a decaying exponential profile, we modified it to generate cross-correlograms with a Gaussian profile, in order to have more synchronous firing for small lags. The Gaussian profile can be seen in Figure 4D, with a standard deviation *σ* = 2.5ms. By setting three different rate levels (5, 10 and 15 Hz) and three different correlation levels (0, 10 and 20 %) this gave rise to 9 conditions, so to 45 recordings in total (5 recordings per conditions, see above).

### 4.3 Template similarity

We define the template for neuron *i* as 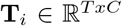, with *T* representing the number of samples and *C* the number of channels. After *flattening* the template by concatenating the signals from different channels 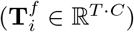, the similarity between two neurons *i* and *j* is quantified via the cosine similarity defined as follows:

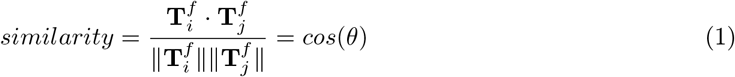

where *θ* is the angle between the two (*T* · *C*)-dimensional vectors 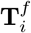 and 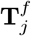. The cosine similarity is therefore bounded between −1 (templates are anti-parallel) and 1 (templates are parallel). A cosine similarity of 0 means that the templates are orthogonal.

### 4.4 Spike sorters

All the spike sorters used in this study were run using the SpikeInterface framework [7], with default parameters. The following are the exact versions that we used for the different spike sorters: Tridesclous (1.6.4), Spyking-circus (1.0.9) [28], Herdingspikes (0.3.7) [13], Kilosort (v1, 2, or 3) [20], YASS (2.0) [15], Ironclust (5.9.8) [8], HDSort (1.0.3) [9]. The desktop machine used has 36 Intel Xeon(R) Gold 5220 CPU @ 2.20GHz, 200Go of RAM and a Quadro RTX 5000 with 16Gb of RAM as a GPU.

### 4.5 Spike sorting comparison

All the quantitative metrics between the results of the spike sorting software and the ground-truth recording were made via the SpikeInterface toolbox.

When comparing a spike sorting output to the ground-truth spiking activity, first an agreement score between each pair of ground-truth and sorted spike trains is computed as:

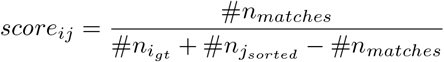

where *#n_i_gt__* and *#nj_sorted_* are the numbers of spikes in the *i*-th ground-truth spike train and the *j*-th sorted spike trains, respectively. *#n_matches_* is the number of spikes within 0.4 ms between the two spike trains.

Once scores for all pairs are computed, an hungarian assignment is used to match ground-truth units to sorted units [7]. All spikes from matched spike trains are then labeled as: true positive (TP), if the spike is found both in the ground-truth and the sorted spike train; false positive (FP), if the spike is found in the sorted spike train, but not in the ground-truth one; and false negative (FN), if the spike is only found in the ground-truth spike train.

After labeling all matched spikes, we can now define these unit-wise performance metrics for each ground-truth unit that has been matched to a sorted unit:

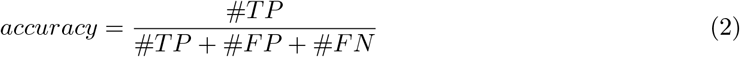

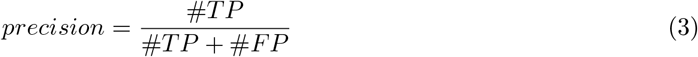

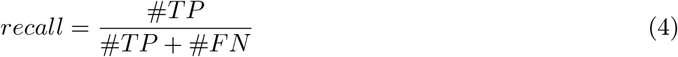

The global accuracy, precision, and recall values shown in Figure 2D are the average values of the performance metrics computed by unit.

Using the unit metrics and the output of the matching procedure, we can further classify each sorted unit as:

**well detected:** sorted units with an accuracy ≥ 0.8
**false positive:** sorted units that are not matched to any ground-truth unit and have a score < 0.2
**redundant:** sorted units that are not the best match to a ground-truth unit but have a score ≥ 0.2
**overmerged:** sorted units with a score ≥ 0.2 with more than one ground-truth unit

In order to generate the spike lag versus recall figures (e.g. Figures 3–5 - figure supplement 1) we expanded the SpikeInterface software with several novel comparison methods and visualization widgets. In particular, we extended the ground-truth comparison class to the `CollisionGTComparison`, which computes performance metrics by spike lag. In addition to the agreement score computation and the matching described in the previous paragraphs, this method first detects and flags all “synchronous spike events” in the ground-truth spike trains. Two spikes from two separate units are considered to be a “synchronous spike event” if their spike times occur within a time lag of 2 ms. The synchronous events are then binned in 11 bins spanning the [−2, 2] ms interval and the *collision recall* is computed for each bin. With a similar principle, we implemented the `CorrelogramGTComparison` to compute the lag-wise relative errors in cross-correlograms between ground-truth units and spike sorted units (Figure 5C).

